# Oxytocin may reduce the accumulation and effects of chronic stress: An exploratory study using hair samples

**DOI:** 10.1101/2025.04.15.649015

**Authors:** Genta Ochi, Nodoka Ohara, Hana Kameo

**Author notes:** **Corresponding author:** Genta Ochi, 1398 Shimami-cho, Kita-ku, Niigata-City, 950-3198, Japan.

## Abstract

**Objective:** This study examined the relationship between hair cortisol concentration (HCC) and hair oxytocin concentration (HOC) and the effects of both hormones on training load and mental health indicators.

**Methods:** Hair samples were collected from 22 female university soccer players, and psychological assessments were conducted in February and March 2024. Cortisol and oxytocin levels were extracted and measured from the hair, and their associations with mood states [measured using the Profile of Mood States 2nd Edition (POMS2)], psychological distress [measured using the Kessler Psychological Distress Scale-6 (K6)], and self-reported training loads were analyzed.

**Results:** Training load and POMS fatigue levels increased significantly from February to March (t = −4.91, P < 0.001, d = 1.52; t = −4.74, P < 0.001, d = 1.01, respectively), but no significant changes were observed in HCC and HOC. HOC significantly moderated the relationship between training load and HCC, with a strong positive correlation between training load and HCC observed in players with “moderate” HOC levels. Additionally, at the second measurement point, a significant negative correlation was observed between HCC and K6 scores, with HOC significantly moderating this relationship. A significant negative correlation was confirmed between changes in POMS vigor and HCC.

**Conclusion:** Combined measurement of HCC and HOC is useful in assessing chronic stress in female soccer players. HOC plays an important role in moderating the relationship between HCC and training load, and mental health indicators. An approach that considers the interactions between these hormones may help prevent overtraining and maintain optimal conditioning.

## Introduction

Athletes who engage in intense daily training to achieve high performance can be in a constant state of significant stress. Insufficient rest, nutrition, or excessive training can cause severe stress, potentially leading to overtraining syndrome characterized by decreased performance and mood (Lehmann et al. 1992; Meeusen et al. 2013). Furthermore, increased risk-taking behaviors due to chronic mental and physical stress may cause injuries (Hughes and Leavey 2012). Understanding stress states may help prevent poor performance, decreased mental health, burnout, and dropout. Many studies and sports settings use questionnaire-based methods to assess stress states; however, psychological indicators can be influenced by events at the time of measurement (such as diet or training), allowing temporary mood to affect the results. Additionally, because the results of mental health and stress measurements may influence evaluations by coaches or managers, athletes may not complete the questionnaires accurately. Therefore, there is a need for physiological assessment methods for chronic stress that are not influenced by temporary mood at the time of measurement.

Recent research has suggested that hormones that accumulate in human hair can serve as useful physiological indicators of long-term stress (Raul et al. 2004). Hair grows approximately 1 cm per month (Wennig 2000), and blood-derived hormones are believed to accumulate via capillaries during hair formation (Gow et al. 2010). Cortisol, commonly used as a stress indicator, accumulates in hair, with elevated levels reported in relation to stressful life events (Dettenborn et al. 2010; Staufenbiel et al. 2015) and during pregnancy (D’Anna-Hernandez et al. 2011). Hair cortisol concentration (HCC) has been found to correlate more strongly with physical stressors, such as intense physical activity, than with psychological stress (Gerber et al. 2012), leading to higher HCC values in individuals with high physical activity levels and athletes who engage in rigorous exercise (Skoluda et al. 2012). This relationship reflects not only inter-individual differences, but also intra-individual variations, as increases in training load among female athletes have been associated with elevated HCC (Sato et al. 2024). However, whether increased chronic stress is associated with a decline in mental health remains unclear. Previous research on hair cortisol has suggested that elevated HCC in athletes may contribute to low mood and reduced aerobic endurance performance (Ochi et al. 2020), but this association with mood decline was not replicated in female athletes. Endocrine responses to stress reactions involve multiple hormones that interact with each other and not cortisol alone. Therefore, the degree of increase in HCC and its effects on mood and performance may be regulated by other hormones that influence individual and group differences.

Oxytocin is known to exert physiological effects that counteract those of cortisol. Oxytocin is a peptide hormone associated with social cognition and attachment (Bielsky and Young 2004), and is secreted during novel events (Ross and Young 2009; Uvnäs-Moberg 1998) and stress responses (Hoehne et al. 2022). Furthermore, oxytocin is involved in stress responses by binding to hypothalamic receptors and exerting inhibitory effects on the HPA axis (Evenepoel et al. 2023; Neumann et al. 2000; Parker et al. 2005). Recent animal studies have reported a positive correlation between hair oxytocin and cortisol (López-Arjona et al. 2021), suggesting that hair oxytocin concentration (HOC) levels may increase under chronic stress conditions. Oxytocin has long been reported to attenuate stress responses (Gibbs 1986) and exert anxiolytic effects (Uvnäs-Moberg and Petersson 2005). This suggests that hair oxytocin may reflect long-term inhibitory effects on the HPA axis. However, many studies have only examined the transient effects of oxytocin and stress, and the relationship between chronic oxytocin levels and stress remains unclear. Similar to cortisol, it is possible to measure the amount of oxytocin accumulated in hair using similar extraction methods, potentially allowing the verification of whether individuals with higher hair oxytocin levels have suppressed cortisol accumulation and reduced negative effects on mood and performance.

Therefore, this study aimed to determine whether oxytocin in hair modulates cortisol secretion and mood effects in female athletes using a two-month longitudinal study with 22 female soccer players.

## Methods

### Participants

Twenty-two members of the women’s soccer club of Niigata University of Health and Welfare consented to participate in this study during the 2024 season (age 19.8 ± 0.8 years, height 160.3 ± 5.5 cm, and weight 54 ± 4.7 kg). For body composition, the players were asked to complete a questionnaire based on the results of their physical examinations received in January.

We provided participants with information regarding the purpose, content, and safety of the study and obtained written consent to participate in the study. The study was conducted in accordance with the principles of the Declaration of Helsinki. The Ethics Committee of the Niigata University of Health and Welfare approved this study (approval number: 18881-220831).

The post-hoc power analysis revealed that with our sample size of 22 participants, a statistical power of 0.80, and a significance level of 0.05 (two-sided), the minimum detectable effect size (*ρ*) was 0.559.

### Experimental procedures

Hair samples were collected, and psychological questionnaires were administered in the second weeks of February and March. January was the off period during the 2024 season, and team training began in February after completion of the off period. The day before the measurement, the participants were given a day off from club activities and were asked to report to the laboratory between 10:00 and 14:00. Participants were asked to refrain from eating or drinking for 2 h before the meeting. During the measurement period, all participants were asked to participate in the activities of their football club. The raters took only two measurements and provided no instructions for any other activity.

### HCC and HOC

Hair samples were collected from the back of the head with minimal variation (Sauvé et al. 2007). To assess chronic stress levels during the previous month, we chose a 1 cm section (10 mg) from the scalp end of the collected hair for analysis, as in previous studies (Greff et al. 2019; Sato et al. 2024). Hair samples were collected using scissors and weighed using an electronic balance (HT84R; Shinko Denshi, Japan). As human hair grows approximately 1 cm per month (Koren et al. 2007; Loussouarn 2001; Wennig 2000), it was assumed that the hair samples from the February and March measurement periods would be completely different. The hair was washed twice with isopropanol for 3 min at room temperature to remove any sweat or sebum secretions adhering to the hair surface, air-dried, and weighed. Washed hair samples were outsourced to Air Plants Bio Co., Ltd. (Japan) for measurement, and the measurement methods were identical to those used in previous studies (Sato et al. 2024).

### Psychological measurements

The participants completed the Profile of Mood States 2nd Edition (POMS2), the Kessler Psychological Distress Scale-6 (K6), and a self-reported training load.

### POMS2

The POMS2 (Yokoyama and Watanabe 2015) comprises 35 items that assess seven mood states (anger– hostility, confusion–bewilderment, depression–ejection, fatigue–inertia, tension–anxiety, vigor–activity, and friendliness). In this study, we used only the 10-item subsets of vigor–activity and fatigue–inertia to minimize the participants’ psychological burden. The internal consistency reliability (Cronbach’s alpha) of this scale in this study was .86 (vigor in February), .92 (vigor in March), .56 (fatigue in February), and .59 (fatigue in March).

### K6

We used the Japanese version of the K6 scale. It is a powerful screening tool for psychological distress (Kessler et al. 2002). Respondents rated six items on a five-point Likert scale, with scores ranging from 0 to 4. A higher total score indicated poorer mental health. Moderate or greater psychological distress was defined as a K6 score of ≥5 (Kessler et al. 2003; Prochaska et al. 2012). The internal consistency reliability (Cronbach’s alpha) of the scale in this study was .79 (February) and .92 (March).

### Self-reported training load

We calculated the self-reported training load as the product of monthly training hours and intensity. We used an 11-point scale ranging from 0 to 10 to measure the rate of perceived exertion (Foster et al. 2001). Self-reported training load, assessed monthly in retrospect, has been validated as having a strong correlation with training volume measured using a heart rate monitor (Matsuura and Ochi 2023). The internal consistency reliability (Cronbach’s alpha) of the scale in this study was .37 (February) and .57 (March).

### Statistical analyses

All analyses were conducted using the R (4.4.2) software (R Foundation, Vienna, Austria). To examine the significance of the changes in each parameter between the two time points (February and March), we used either the Wilcoxon signed-rank test or the paired t-test, depending on data normality. Additionally, to investigate the relationships between training load, HCC, and HOC, we used Spearman’s rank correlation coefficient (ρ). This metric does not assume a normal distribution, making it suitable for analyzing physiological data. Furthermore, to examine the effect of training load on HCC and the moderating effect of HOC on this relationship in detail, we conducted interaction analysis using linear regression models. To better understand the effects of HOC, we performed a subgroup analysis by dividing the HOC values into tertiles (low, medium, and high). For associations between changes, we calculated the differences between two time points for each variable and examined their relationships using Spearman’s correlation and regression analyses. We also analyzed the moderating effect of HOC on the relationship between psychological indicators (K6, POMS fatigue, and POMS vigor) and HCC using a similar method. For all statistical tests, the significance level was set at less than 5%, and statistical values and effect sizes were calculated.

## Results

### Overview

The main parameters measured in this study are as follows: training load increased significantly from the first to the second measurement (t = −4.91, P < 0.001, d = 1.52), as did POMS fatigue (t = −4.74, P < 0.001, d = 1.01). In contrast, no significant difference was observed in HCC between the first and second measurements (V = 115, P = 0.73). Regarding HOC, no significant change was observed between the first and second measurements (t = −0.89, P = 0.38). The K6 scores showed an increasing trend from the first to the second measurement, but did not reach statistical significance (V = 32.5, P = 0.12). For POMS vigor, no significant difference was observed between the first and second measurements (t = −0.60, P = 0.55). For the cognitive function indicators, Stroop interference effects (horizontal and vertical tasks), no significant changes were observed between the first and second measurements (P = 0.94, P = 0.50, respectively).

### Relationship between training load, HCC, and HOC

Table 2 presents the correlation coefficients for each variable. No significant correlations were observed between HOC and HCC during either the February (ρ = −0.03, P = 0.90) or the March (ρ = −0.24, P = 0.28) measurements. At both measurements, no correlation was observed between training load and HCC (February: ρ = 0.35, P = 0.11; March: ρ = −0.18, P = 0.41). No correlation was found between February HCC and any psychological scales, but March HCC showed a negative correlation with K6 (ρ = −0.65, P < 0.05). The change in HCC from February to March showed a negative correlation with POMS vigor (ρ = −0.44, P < 0.05), and the change in HOC showed a positive correlation with K6 (ρ = 0.43, P < 0.05).

**Table 1.** Changes in main variables between February and March measurements.

| Variable | February |  | March |  | Statistic | Type | P Value |
| --- | --- | --- | --- | --- | --- | --- | --- |
|  | Mean | SD | Mean | SD |  |  |  |
| Training Load | 1207.09 | 967.95 | 3466.82 | 1788.12 | -4.91 | t-test | <0.001 |
| K6 | 8.64 | 3.36 | 10.23 | 4.53 | 32.5 | Wilcoxon | 0.12 |
| POMS Fatigue | 2.36 | 1.79 | 4.59 | 2.58 | -4.74 | t-test | <0.001 |
| POMS Vigor | 6.41 | 3.98 | 6.95 | 3.99 | -0.6 | t-test | 0.55 |
| Cortisol (HCC) | 10.27 | 5.69 | 9.76 | 3.07 | 115 | Wilcoxon | 0.73 |
| Oxytocin (HOC) | 3.57 | 1.03 | 3.83 | 1.3 | -0.89 | t-test | 0.38 |

**Table 2.** Correlation matrix.

|  |  | K6 | POMS Fatigue | POMS Vigor | Training Load | HCC | HOC |
| --- | --- | --- | --- | --- | --- | --- | --- |
| Feb | K6 |  | 0.45* | 0.13 | 0.28 | -0.04 | -0.02 |
|  | POMS Fatigue | 0.45* |  | 0.4 | -0.01 | 0.14 | -0.05 |
|  | POMS Vigor | 0.13 | 0.4 |  | -0.01 | -0.24 | 0.37 |
|  | Training Load | 0.28 | -0.01 | -0.01 |  | 0.35 | 0.03 |
|  | HCC | -0.04 | 0.14 | -0.24 | 0.35 |  | -0.03 |
|  | HOC | -0.02 | -0.05 | 0.37 | 0.03 | -0.03 |  |
| Mar | K6 |  | 0.49* | 0.06 | -0.08 | -0.65* | 0.06 |
|  | POMS Fatigue | 0.49* |  | 0.11 | 0.1 | -0.31 | 0.23 |
|  | POMS Vigor | 0.06 | 0.11 |  | 0.02 | 0.07 | -0.03 |
|  | Training Load | -0.08 | 0.1 | 0.02 |  | -0.18 | 0.17 |
|  | HCC | -0.65* | -0.31 | 0.07 | -0.18 |  | -0.24 |

|  | HOC | 0.06 | 0.23 | -0.03 | 0.17 | -0.24 |  |
| --- | --- | --- | --- | --- | --- | --- | --- |
| Change | K6 |  | 0.43* | -0.22 | 0.33 | -0.03 | 0.43* |
|  | POMS Fatigue | 0.43* |  | -0.07 | 0.44* | 0.2 | 0.25 |
|  | POMS Vigor | -0.22 | -0.07 |  | 0.15 | -0.44* | -0.03 |
|  | Training Load | 0.33 | 0.44* | 0.15 |  | 0.03 | 0.37 |
|  | HCC | -0.03 | 0.2 | -0.44* | 0.03 |  | 0.28 |
|  | HOC | 0.43* | 0.25 | -0.03 | 0.37 | 0.28 |  |

*Note.* The correlation coefficients (Spearman's rho) between each variable are shown (\* $P < 0.05$ ). Feb, February; Mar, March.

Notably, linear regression analysis confirmed a significant moderating effect of HOC on the relationship between training load and HCC at the February measurement (training load × HOC: β = −0.0029, P = 0.025). In this interaction model, both training load (β = 0.012, P = 0.013) and HOC (β = 3.95, P = 0.034) showed significant positive main effects on HCC. In the subgroup analysis by HOC level, a strong positive correlation between training load and HCC was observed particularly in the “medium” level HOC group (ρ = 0.61, P = 0.026). In contrast, this association was not significant in the “low” and “high” HOC groups (Figure 1). Analysis of the three-factor interaction model (HOC × training load × time point) revealed a significant interaction among the three factors (β = 0.0034, P = 0.004). This indicates that the moderating effect of HOC on the relationship between training load and HCC differed by measurement point.

**Fig 1.**
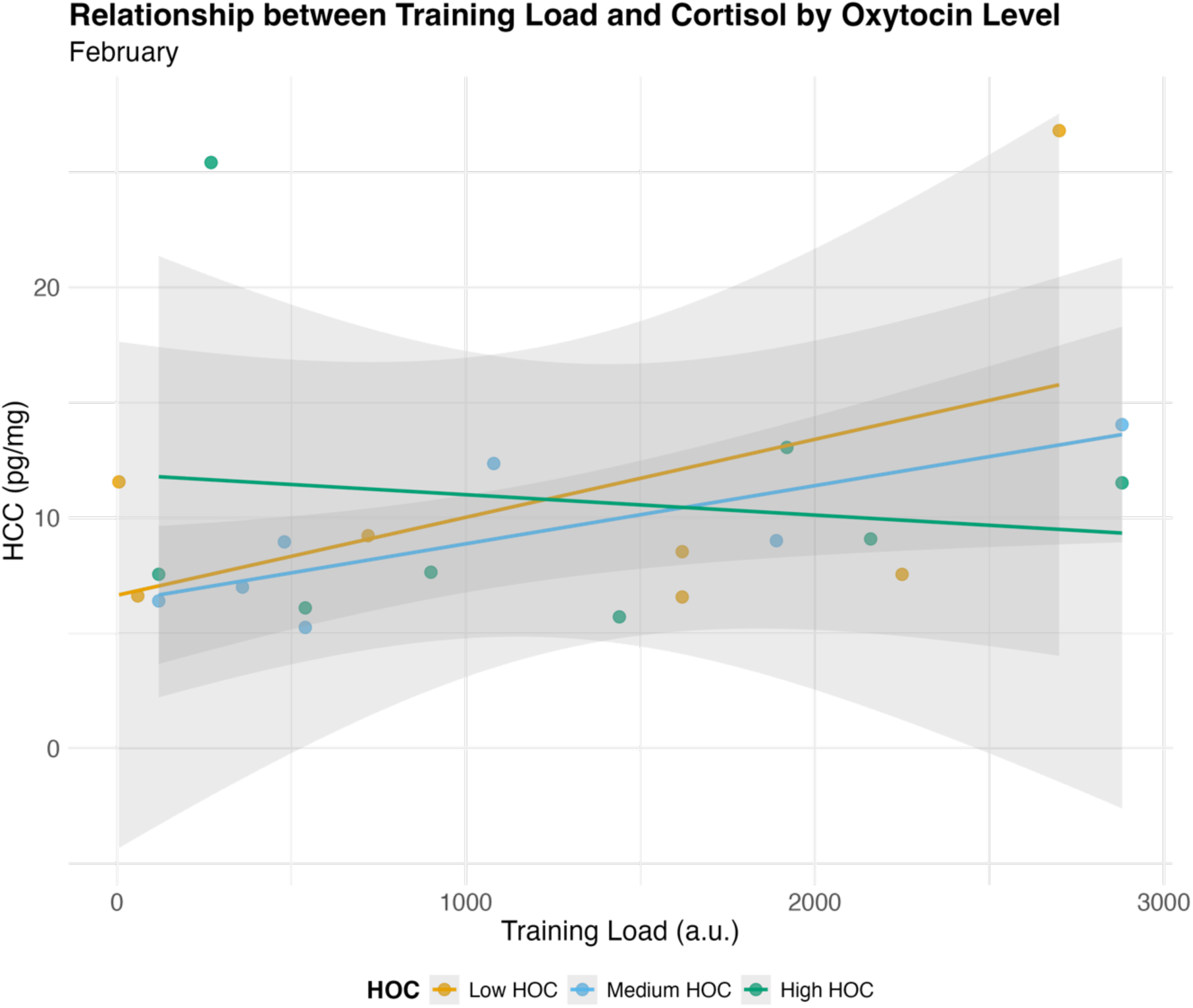
Relationship between training load and HCC by HOC level in February

### Role of HOC in the relationship between HCC and psychological indicators

At the second measurement, a significant negative correlation was observed between HCC and K6 scores (ρ = −0.65, P = 0.001). Additionally, a significant moderating effect of HOC was confirmed in the relationship between HCC and K6 (β = −0.67, P = 0.006). Regarding POMS fatigue, although no significant correlation was observed with HCC (ρ = −0.31, P = 0.17), a significant moderating effect of HOC was observed in the relationship between HCC and POMS fatigue (β = −0.51, P < 0.001). A significant negative correlation was observed between the changes in POMS vigor and HCC (ρ = −0.44, P = 0.039)(Figure 2). However, the moderating effect of HOC changes on this relationship was not significant. No significant correlations were observed between HCC and other psychological or cognitive function indicators at any time point.

**Fig 2.**
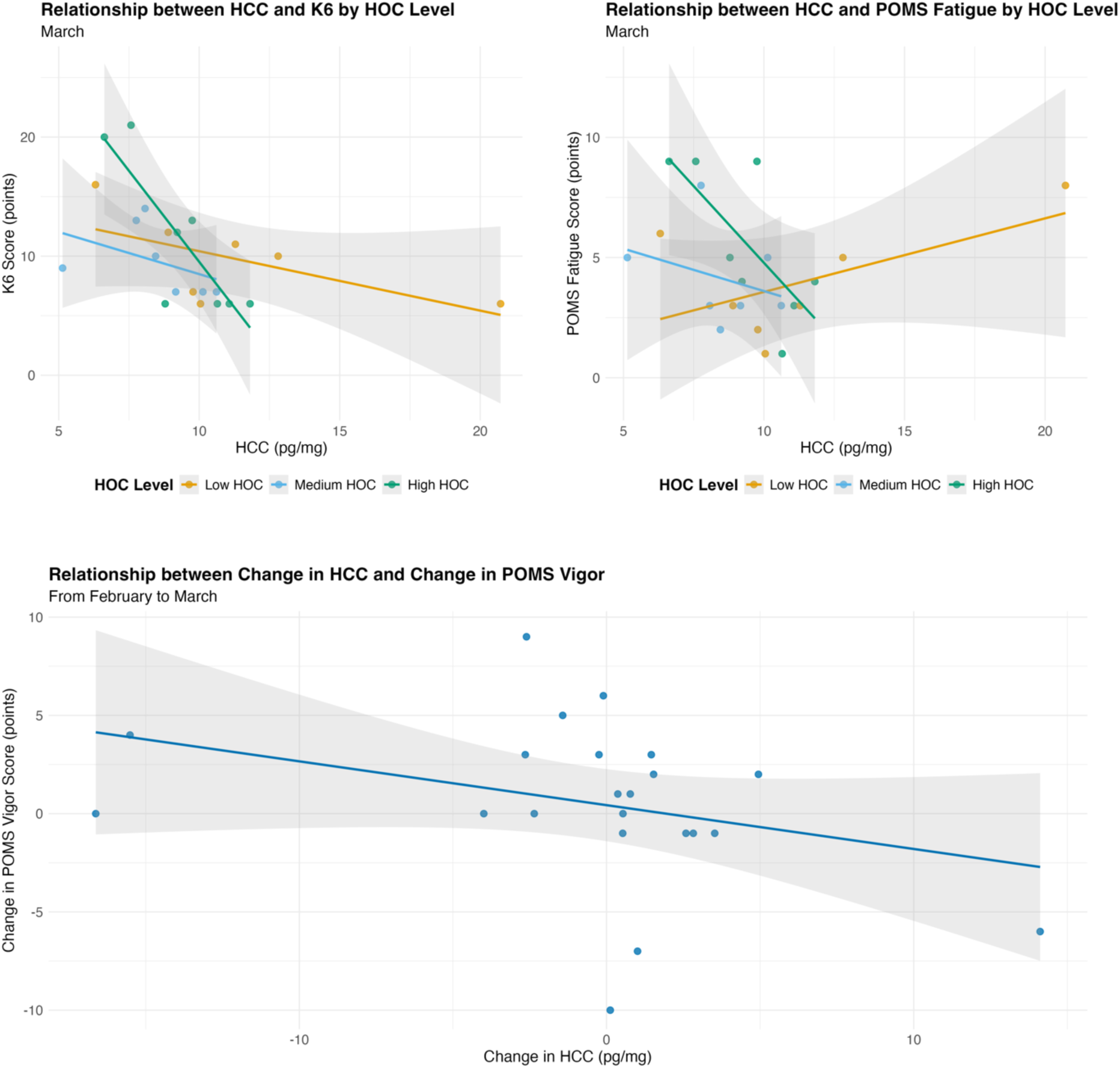
Relationship between mental health and HCC by HOC level

### Analysis of changes

No significant correlation was observed between the changes in training load and HCC (ρ = 0.03, P = 0.89). Similarly, no clear association was observed between changes in HOC and HCC (ρ = 0.28, P = 0.21). In the partial correlation analysis controlling for changes in the training load, the association between HOC and HCC changes was not significant (ρ = 0.29, P = 0.21). In the regression analysis of changes (HCC change ∼ training load change × HOC change), the interaction effect was not statistically significant (β = 0.0008, P = 0.25). These results indicate that there are complex relationships between training load, HCC, HOC, and psychological indicators, with HOC having a moderating effect on the relationship between training load and HCC, particularly in the first measurement. Additionally, in the second measurement, HOC was found to have a moderating effect on the relationship between HCC and psychological indicators, especially K6 and POMS fatigue.

## Discussion

In this study, we measured HCC and HOC to evaluate chronic stress states in female soccer players, and examined their relationships with training load, mood indicators, and cognitive function. The measurements were taken at two time points: February and March. The training load and POMS fatigue level increased significantly, but no significant changes were observed in the HCC and HOC. Notably, HOC was confirmed to have a moderating effect on the relationship between training load and HCC, with a strong positive correlation between training load and HCC particularly observed in players with “moderate” HOC levels.

Among the parameters measured in this study, training load and POMS fatigue levels increased significantly from February to March. This likely reflects an increase in the physical load associated with the start of team training following the off-season. Although the K6 scores showed an upward trend, they did not reach statistical significance. These results suggest that although players experienced physical fatigue due to increased training intensity, the notable adverse effects on their mental health were limited.

Interestingly, although a previous study (Sato et al. 2024) found that an increased training load was associated with elevated HCC, this study observed no significant changes in HCC between the two time points. This discrepancy in results may be explained by the moderating effect of HOC. Specifically, at the first measurement point, training load and HOC showed significant positive main effects on HCC, whereas the interaction between training load and HOC demonstrated a significant negative effect. This indicates that the impact of the training load on HCC varies depending on the HOC level. Subgroup analysis revealed a strong positive correlation between training load and HCC in the “moderate” HOC group, but this association was not significant in the “low” and “high” HOC groups. These findings suggest that oxytocin plays a complex role in stress response, which is consistent with the inhibitory effects of oxytocin on the HPA axis reported in previous studies (Evenepoel et al. 2023; Neumann et al. 2000; Parker et al. 2005).

The moderating effect of HOC on the relationship between HCC and psychological indicators is noteworthy. At the second measurement point, a significant negative correlation was observed between HCC and K6 scores, and HOC significantly moderated this relationship. Similarly, a significant moderating effect of HOC was confirmed in the relationship between HCC and POMS fatigue levels. These results suggest that the psychological impact of HCC differs depending on HOC levels. Specifically, at certain HOC levels, a lower HCC may be paradoxically associated with mental stress, reflecting a complex interaction between oxytocin and cortisol.

Additionally, a significant negative correlation was observed between changes in POMS vigor and HCC. This finding suggests that a decrease in HCC is associated with increased vigor. However, the moderating effect of HOC change on this relationship was not significant. This result partially aligns with a previous study (Ochi et al. 2020) that reported an association between HCC and vigor but contrasts with findings that this association was not confirmed in female athletes. The results of this study suggest a relationship between HCC and mood states in female soccer players. However, this relationship may be moderated by HOC.

In the analysis of change values, no significant correlation was observed between changes in training load and HCC. Similarly, no clear association was found between the changes in HOC and HCC. These results indicate that the relationships among training load, HCC, HOC, and psychological indicators are complex, and that interactions at specific time points are important.

## Limitation

This study had several limitations. First, we did not measure cortisol in the blood or saliva simultaneously with hair sampling, making it impossible to examine the relationship between transient cortisol and oxytocin levels, and HCC and HOC effects. A previous research (Ochi et al. 2020) reported no correlation between HCC and blood cortisol concentration. Additionally, we confirmed that salivary cortisol and oxytocin levels at the time of measurement did not correlate with HCC or HOC, indicating that they are independent indicators (preliminary experimental data). Therefore, it is likely that the HCC and HOC measurements in this study were not influenced by transient stress at the time of measurement. Although this study targeted all players belonging to a single university’s women’s soccer team, the measurement period occurred when senior players had retired and before new students joined, resulting in a limited sample size. Furthermore, although this study focused on players from a women’s soccer team with relatively similar lifestyle habits and stressors, it remains unclear whether these results can be applied to other women’s soccer teams or to male players. Questions remain regarding the sex-based differences in HCC (Raul et al. 2004; Schell et al. 2017), and the impact of HCC on mental health has been reported to be stronger in women than in men (Kim et al. 2021). Future studies should conduct analyses with more participants and verification in male soccer players. Finally, regarding the mechanism of interaction between oxytocin and cortisol, it is difficult to infer causal relationships from the observational design of this study. Future research should employ experimental approaches to elucidate these hormonal interactions in more detail as well as approaches involving long-term measurements and analyses.

## Conclusion

The results of this study demonstrated that the relationship between training load and HCC is moderated by HOC, with a strong positive correlation between training load and HCC particularly observed in players with “moderate” HOC levels. Additionally, the moderating effects of HOC were observed in the relationship between HCC, K6 scores, and POMS fatigue levels. These findings indicated that oxytocin and cortisol play important roles in the assessment of chronic stress in female athletes. Specifically, the combined measurement of HCC and HOC may be useful for a more accurate evaluation of the accumulation of physical stress from training and its psychological impacts. This measurement approach has the potential to function as a physiological indicator that visualizes the stress states of individual players and contributes to the early detection of burnout and overtraining, thereby providing a new perspective for athlete health management and performance enhancement.

## Acknowledgments

We would also like to thank Editage (www.editage.com) for English language editing.

## Statements and declarations

### Ethical Considerations

We provided the participants with information regarding the purpose, content, and safety of the study, and they provided written consent to participate. The study was conducted in accordance with the principles of the Declaration of Helsinki. The Ethics Committee of the Niigata University of Health and Welfare approved this study (approval number: 18881-220831).

### Data Availability

The data that support the findings of this study are available from the corresponding author upon reasonable request.

### Author contributions

GO conceptualized the research, investigated the data, conducted formal analysis, and wrote the original draft, and reviewed and edited the final version. NO and HK investigated the data and conducted formal analysis. All authors have read and approved the manuscript.

## Abbreviations

HCC: hair cortisol concentration
HOC: hair oxytocin concentration
HPA: 
K6: Kessler Psychological Distress Scale-6
POMS2: Profile of Mood States 2nd Edition

